# Pyrethroid Resistance Intensity of *Anopheles gambiae sensu lato* (Diptera: Culicidae) from Phase II Hut Trial Station in KOLOKOPE, Eastern Plateau Togo: A Potential Site to Assess the Next Generation of Long-Lasting Insecticidal Nets

**DOI:** 10.1101/2020.08.19.257519

**Authors:** Koffi Mensah Ahadji-Dabla, Joseph Chabi, Georges Apétogbo, Edoh Koffi, Melinda Patricia Hadi, Guillaume Koffivi Ketoh

## Abstract

Per WHO recommendations, the implementation of the next-generation of Long Lasting Insecticidal Nets (LLINs) for malaria vector control requires appropriate investigations on the insecticide resistance profile of the vector and the impact of the LLINs on the known resistant mosquitoes. The next-generation of LLINs are actually an incorporation of a mixture of pyrethroid insecticides and a synergist such as PBO. Several studies have proven the additional impact of PBO on the increase in the mortality rate of *Anopheles gambiae* s.l. (Diptera: Culicidae). However, further assessments need to be done at community level in order to set a stage for the acceleration of the WHO policies on the implementation of the next-generation of LLINs. Kolokopé is a cotton-growing area in the central region of Togo characterized by an intensive use of agro pesticides and insecticides. A phase II experimental hut station for the evaluation of mosquito control tools has been built in Kolokopé. For the characterization of the site, WHO susceptibility tests using diagnostic doses of eight insecticides, PBO synergist assay and intensity assay of three pyrethroids (5x and 10x) were conducted on adult female mosquitoes obtained from larvae collected around the site. *Anopheles gambiae* s.l. from Kolokopé showed high resistance to pyrethroids and DDT, but in lesser extent to carbamates and organophosphates. Likewise, high intensity of resistance to pyrethroid was observed with less than 40% mortality at 10x deltamethrin, 52 and 29% mortality at 10x permethrin and 10x alphacypermethrin, respectively. Also, the addition of PBO showed a reversal mortality which was similar to mortality rate at 10x doses of pyrethroids. The high pyrethroid intensity resistance recorded at Kolokopé could be mainly due to the pressure on *An. gambiae s.l*. through the excessive use of insecticide in agriculture. This can be used for the assessment of the next-generation of LLINs either in experimental hut or at a community trial.

## Introduction

The spread of insecticide resistance in malaria vectors has been a source of constant concerns for malaria endemic countries worldwide. In Sub Sahara African countries, insecticide resistance in *Anopheles gambiae s.l*. (*An. gambiae*) has been worsen by the massive distribution of Long Lasting Insecticidal Nets (LLINs) and Indoor Residual Spraying (IRS) campaigns. The use of pyrethroids through different application methods, puts considerable resistance selection pressure on many pests of importance in public health, particularly on malaria vectors (Nauen 2007). Data from the latest malaria report of the 76 malaria endemic countries that reported standard monitoring data for 2010 to 2016, show that resistance was detected in 61 (54%) countries to at least one insecticide in one malaria vector from a collection site. The same report stated that from 2010 to 2016, malaria endemic countries that reported pyrethroid resistance increased from 71 to 81% (WHO 2017).

Pyrethroid insecticides have been used since 1980’s and the resistance to this class of insecticide has been reported in many countries, especially in West Africa (Elissa et al., 1993, Chandre et al. 1999, Diabaté et al. 2002, Ahadji-Dabla et al. 2014, Chabi et al. 2016, Ketoh et al. 2018), and it has affected vector control strategies based on the use of LLINs. In these countries, there is a need to develop and deploy new tools, as stated in the third pillar of the Global Plan for Insecticide Resistance Management in Malaria Vectors (GPIRM), launched in 2012 by the World Health Organization (WHO) (WHO 2012). Malaria vectors developed several mechanisms allowing them to be less susceptible to pyrethroids. These include *kdr* mutation (*kdr* L1014F in West Africa and *kdr* L1014S in East Africa) and over-transcription of detoxification enzymes. Hence, in some areas, malaria vector populations subjected to the diagnostic doses show more and more decrease in susceptibility to pyrethroids and many other insecticides. Therefore, the need to develop new molecules or combination of insecticides to control resistant mosquitoes has become urgent. As part of the new molecules and tools, Actellic 300CS, a reformulated organophosphate insecticide has be developed by Syngenta in collaboration with Innovative Vector Control Consortium (IVCC) (Rowland et al. 2013). Alphacypermethrin SB an IRS insecticide, LLINs such as Akanet LN, Miranet LN, Olyset Duo LN, Panda Net 2.0 LN, Veeralin LN, Yahe LN, and a larvicide, VectoMax GR, are under field testing (Hemingway et al. 2016). However, the effectiveness of the new molecules for IRS and tools (LLINs) in controlling resistance in mosquitoes need to be well assessed in appropriate sites and countries using standard procedures such as the Phase II hut trial.

This study was therefore initiated to characterize and profile the *An. gambiae s.l*. mosquitoes from Kolokopé, where about seven standard experimental huts have been set up to evaluate new vector control tools.

## Material and Methods

### Study site

Kolokopé is a village located in the Plateau region of Togo (07°47′59″N, 01°18′E), about 200 km from Lomé, the capital city. The region is characterised by a long rainy season from March to October and a dry season from November to February. The annual rainfall is estimated at 1300-1500mm per year. Farming activities are the main source of revenue of the population and specifically cotton cultivation with approximately 236 hectares of land and an estimated production of 1000 tons per year. Huge quantity of insecticides are used in the area for crop protection (CRA-SH 2012), and resistance to deltamethrin and permethrin has been recently reported in the area (Ketoh et al. 2018).

### WHO Susceptibility Test

Susceptibility testing of *An. gambiae s.l*. populations from Kolokopé was done using the WHO test kits according to standard testing protocols (WHO 2013). Four classes of insecticides, including (1) pyrethroids (0.05% deltamethrin, 0.75% permethrin, and 0.05% lambdacyhalothrin), (2) organochlorine (4% DDT), (3) organophosphates (1% fenitrothion, 5% malathion, and 0.4% chlorpyrifos methyl), and (4) carbamates (0.1% propoxur and 0.1% bendiocarb) were tested at the diagnostic concentrations (DC). Additionally, synergist assays were performed using piperonyl butoxide synergist (4% PBO) and intensity assays using 5x and 10x diagnostic doses of deltamethrin, permethrin, and alphacypermethrin (WHO 2016). Regarding intensity assays, 98-100% mortality rates at 5×DC indicates that there is no need for assay at the 10×DC (low resistance intensity); mortality rates less than 98% at the 5×DC (moderate resistance intensity, 10×DC assay needed); 98-100% mortality rates at the 10×DC confirms a moderate resistance intensity; and finally, high resistance intensity is indicated by mortality rates less than 98% at 10×DC.

*Anopheles gambiae s.l*. larvae were collected around the study site in October 2017, brought to the field insectary and reared to adulthood. Twenty to twenty-five non-blood fed females aged 3-5 days were exposed for one hour to different doses of above-mentioned insecticides. For the synergist assay, mosquitoes were pre-exposed to PBO for one hour before exposure to the insecticide for an additional hour. The number of mosquitoes knocked down was recorded every 10 minutes for 1 hour, and mortality recorded after 24 hours (WHO 2016). Tests with silicone and olive oil impregnated papers were run in parallel and served as controls. The susceptible *An. gambiae*, Kisumu strain was used as reference.

### Species identification and *kdr* L1014F and *ace1* G119S detection

*Anopheles* specimens from the susceptibility testing stored at −20°C, were randomly selected for PCR analyses. SINE-PCR was used for species identification (Santolamazza et al. 2008). The detection of *kdr* L1014F and *ace1* G119S alleles was conducted following the methods of Martinez-Torres et al. (1998) and Weill et al. (2004), respectively.

DNA extraction was done on a whole mosquito using the protocol designed by Collins et al. (1987) and amplified in 20μl of a master mixture. Specifically, (5’-TCG-CCT TAG ACC TTG CGT TA-3’) and (5’-CGC TTC AAG AAT TCG AGA TAC-3’) are the primers used. With regard to the amplification conditions, activation of the DNA polymerase was the initial step done for 10min at 94°C followed by 35 cycles at 94°C for 30s. Hybridization was done at 54°C for 30s and at 72°C for 1min, respectively, and finally, elongation was done at 72°C for 10min followed by a decrease in temperature to 4°C. All amplified fragments were analyzed by electrophoresis on 2% agarose gel and visualized under UV light after being stained with ethidium bromide.

The detection *kdr* L1014F mutation was performed with: common primers, Agd1 (5’-ATA GAT TCC CCG ACC ATG-3’) and Agd2 (5’-AGA CAA GGA TGA TGA ACC-3’), susceptible primer, Agd3 (5’-AAT TTG CAT TAC TTA CGA CA-3’), and resistance primer, Agd4 (5’-CTG TAG TGA TAG GAA ATT TA-3’). After DNA polymerase activation (94°C for 3s followed by 35 cycles at 94°C for 30s), followed the hybridization (30s at 55°C and 10s at 72°C) and finally the elongation (5min at 72°C).

To detect the presence of the *ace1* G119S mutation, a PCR-Restriction Fragment Length Polymorphism (PCR-RFLP). A master mixture of 25 μl PCR reaction of 1 μl each of 10 μM Primers EX3AGdir (GATCGTGGACACCGTGTTCG) and EX3AGrev (AGGATGGCCCGCTGGAACAG), 12.5 μl of GoTaq, 9 μl of DNase-free water and 1.5 μl of 1/40 dilution of DNA template was prepared. An enzymatic digestion step was followed after the PCR reaction. A 20 μl restriction enzyme reaction mixture of 2 μl of Enzymatic Buffer B 10X, 0.2 μl of Acetylated BSA at 10 μg/μl and 0.5 μl of 10 U/μl restriction enzyme Alu I (Promega), 12.3 μl of DNase-free water and 5 μl of PCR products, was prepared. Incubated was done at 37 °C for 4 h in a thermocycler. The resulting products were run on 2 % agarose gels stained with ethidium bromide and visualized under UV light.

## Results

### Resistance status of *Anopheles gambiae* s.l

According to the standard WHO criteria, the wild population of *An. gambiae s.l*. tested is highly resistant to deltamethrin, permethrin and alphacypermethrin at 10xDC, with mortality rates of 39, 52, and 29%, respectively. The mortality rates increase when mosquitoes were exposed to PBO before being exposed to DCs of deltamethrin and permethrin. Mosquitoes were resistant to the DC of all other insecticides tested expect to malathion, where a suspected resistance was observed (96.6%).

### *Anopheles* species and *kdr* L1014F and *ace1* G119S mutations associated

Two species (n = 176) were identified at Kolokopé (see Table 2). *Anopheles gambiae* was more frequent both in WHO susceptibility testing and experimental huts samples with 98.8 and 97.9%, respectively. *Anopheles coluzzii* represented less than 1%. The knockdown *L1014F* allele was present at high frequency (> 0.9) especially in *An. gambiae* whereas *G119S* allele was present at low frequency in both species.

**Table 1:**
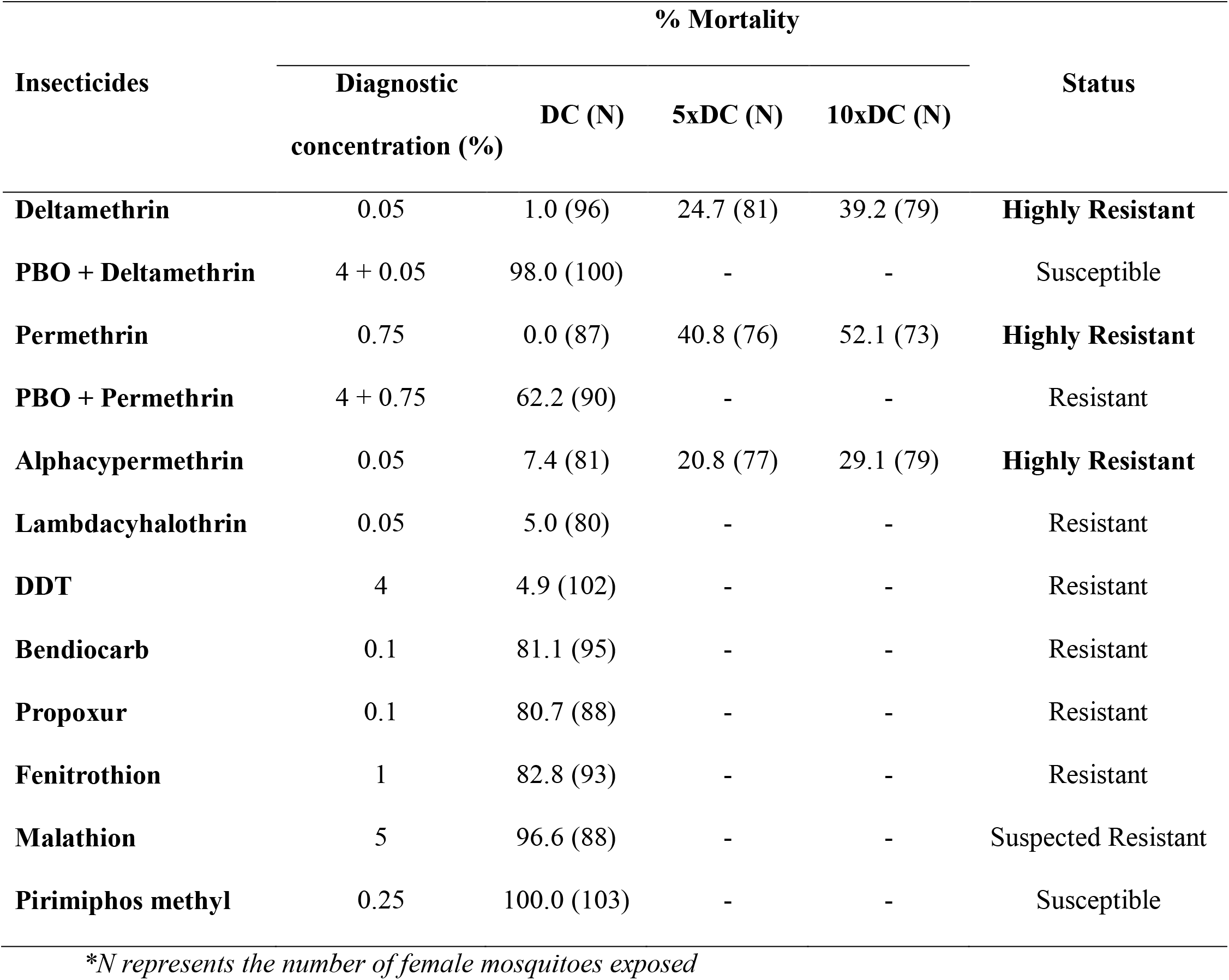
Resistance profile of *An. gambiae s.l*. from Kolokopé

| Insecticides | % Mortality |  |  |  | Status |
| --- | --- | --- | --- | --- | --- |
|  | Diagnostic<br>concentration (%) | DC (N) | 5xDC (N) | 10xDC (N) |  |
| **Deltamethrin** | 0.05 | 1.0 (96) | 24.7 (81) | 39.2 (79) | **Highly Resistant** |
| **PBO + Deltamethrin** | 4 + 0.05 | 98.0 (100) | - | - | Susceptible |
| **Permethrin** | 0.75 | 0.0 (87) | 40.8 (76) | 52.1 (73) | **Highly Resistant** |
| **PBO + Permethrin** | 4 + 0.75 | 62.2 (90) | - | - | Resistant |
| **Alphacypermethrin** | 0.05 | 7.4 (81) | 20.8 (77) | 29.1 (79) | **Highly Resistant** |
| **Lambdacyhalothrin** | 0.05 | 5.0 (80) | - | - | Resistant |
| **DDT** | 4 | 4.9 (102) | - | - | Resistant |
| **Bendiocarb** | 0.1 | 81.1 (95) | - | - | Resistant |
| **Propoxur** | 0.1 | 80.7 (88) | - | - | Resistant |
| **Fenitrothion** | 1 | 82.8 (93) | - | - | Resistant |
| **Malathion** | 5 | 96.6 (88) | - | - | Suspected Resistant |
| **Pirimiphos methyl** | 0.25 | 100.0 (103) | - | - | Susceptible |

\*N represents the number of female mosquitoes exposed

**Table 2:** Species composition and resistance mechanisms of *An. gambiae s.l*. from Kolokope sampled from two different collection methods

| Type of sample | Species | Total | KdrW GENOTYPING |  |  |  |  | ACE-1 GENOTYPING |  |  |  |  |
| --- | --- | --- | --- | --- | --- | --- | --- | --- | --- | --- | --- | --- |
|  |  |  | RR | RS | SS | Total | FREQUENCY | RR | RS | SS | Total | FREQUENCY |
| **WHO**<br>**susceptibility**<br>**test** | *Anopheles gambiae s.s* | 87 (98.8%) | 86 | 0 | 1 | 87 | 0.99 | 1 | 19 | 60 | 80 | 0.13 |
|  | *Anopheles coluzzii* | 1 (0.12%) | 1 | 0 | 0 | 1 | 1.00 | 0 | 0 | 1 | 1 | 0.00 |
|  | **Total** | **88** | **87** | **0** | **1** | **88** | **0.99** | **1** | **19** | **61** | **82** | **0.13** |
| **Experimental**<br>**hut** | *Anopheles gambiae s.s* | 86 (97.7%) | 83 | 1 | 2 | 86 | 0.97 | 0 | 17 | 67 | 84 | 0.10 |
|  | *Anopheles coluzzii* | 2 (0.23%) | 0 | 1 | 1 | 2 | 0.25 | 0 | 1 | 1 | 2 | 0.25 |
|  | **Total** | **88** | **83** | **2** | **3** | **88** | **0.95** | **0** | **18** | **68** | **86** | **0.10** |

## Discussion

According to the Insecticide Resistance Action Committee, resistance is “a heritable change in the sensitivity of a pest population that is reflected in the repeated failure of a product to achieve the expected level of control when used according to the label recommendation for that pest species” (IRAC 2018). Based on the mortality rates recorded at the 10×DC, this study showed that the resistance status of *An. gambiae* s.l. to pyrethroids was far beyond the WHO recommended diagnostic doses. Cotton-growing is the main agriculture practice at Kolokopé with an estimated production of 1000 tons per year; such practice requires the use of a significant amount of insecticides and fertilizers in the area. A study by Yadouleton et al. (2011) reported a regular use of fertilizers and insecticides in cotton fields in Benin.

Insecticide resistance monitoring activities of *An. gambiae* s.l. have been conducted in several areas of Togo and showed high level of pyrethroid resistance at Kolokopé (Ketoh et al. 2018, Amoudji et al. 2019) or surroundings (Djègbè et al. 2018). This study reports high pyrethroid intensity resistance in *An. gambiae s.l*. at Kolokopé; similar results were recently reported in Ghana (Pwalia et al. 2019), Mali (Sovi et al. under review), and Nigeria (Awolola et al. 2018) and were attributed to the selection pressure on vector populations following the rapid scale up and use of pyrethroid-based vector control interventions and pyrethroid insecticides use in agriculture.

To date in Togo, malaria vector control relies exclusively on the use of LLINs, one of the strategies recommended by the WHO (WHO 2018). Therefore, the National Malaria Control Program of Togo had distributed a total of 4,706,417 LLINs in 2017, through national and routine campaigns. The bad news in this study is the high level of resistance, sounding the bell of an urgent implementation of insecticide resistance management program. Nevertheless, the good news is that Kolokopé becomes an area par excellence where the efficacy of new malaria vector control tools could be evaluated in experimental huts constructed for that purpose. We can hypothesized that any tool that could control efficiently the *Anopheles* strain of Kolokopé, could also control at least the other wild strains across the country.

The susceptibility tests clearly showed that when the mosquitoes were exposed to PBO prior to pyrethroids, the mortality was reversed. Experimental hut trials conducted at Kokolopé in 2013, showed the efficacy of two LLINs: PermaNet^®^ 3.0 and Olyset^®^ Plus. These LLINs are PBO+deltamethrin and PBO+permethrin incorporated, respectively (Ketoh et al. 2018). Also, a study by Dadzie et al. (2017) in Ghana, reported the role of PBO in enhancing the efficacy of pyrethroid insecticides against *An. gambiae s.l*.

It is important to emphasize the fact that two species (*An. gambiae* and *An. coluzzii*) were identified in this study with *An. gambiae* commonly represented and having a high *kdr* mutation frequency. However, it was previously reported that each of both species was almost ~50% frequent and *An. coluzzii* had the highest *kdr* mutation frequency. This situation could be explained by the dynamics of species composition and resistance allele frequency (Stump et al. 2004, Chouaïbou et al. 2008, Djègbè et al. 2011).

## Conclusion

This study reveals the high pyrethroid intensity resistance recorded at Kolokopé which could be mainly due to the pressure on *An. gambiae s.l*. through the excessive use of insecticide in agriculture. Piperonyl butoxide reversed pyrethroids mortality rates. This can be used for the assessment of the next-generation of LLINs either in experimental hut or at a community trial.

## Acknowledgments

Authors would like to thank to local community of Kolokopé. K. M. Ahadji-Dabla thanks the Fulbright Visiting Scholar Program. All authors read and approved the final version of the manuscript. The authors declare that they have no competing interests.

## References

Ahadji-Dabla, K. M., G. K. Ketoh, W. S. Nyamador, G. Y. Apétogbo, and I. A. Glitho. 2014. Susceptibility to DDT and pyrethroids, and detection of knockdown resistance mutation in *Anopheles gambiae sensu lato* in southern Togo. Int. J. Biol. Chem. Sci. 8: 314–323.

Amoudji, A. D., K. M. Ahadji-Dabla, A. S. Hien, Y. G. Apétogbo, B. Yaméogo, D. D. Soma, R. Bamogo, R. T. Atcha-Oubou, R. K. Dabiré, and G. K. Ketoh. 2019. Insecticide resistance profiles of *Anopheles gambiae s.l.* in Togo and genetic mechanisms involved, during 3-year survey: is there any need for resistance management? Malar. J. 18: 177.

Awolola, T. S., A. Adeogun, A. K. Olakiigbe, T. Oyeniyi, Y. A. Olukosi, H. Okoh, T. Arowolo, J. Akila, A. Oduola, C. N. Amajoh. 2018. Pyrethroids resistance intensity and resistance mechanisms in *Anopheles gambiae* from malaria vector surveillance sites in Nigeria. PLoS ONE. 13(12): e0205230.

Chabi, J., P. K. Baidoo, A. K. Datsomor, D. Okyere, A. Ablorde, A. Iddrisu, M. D. Wilson, S. K. Dadzie, H. P. Jamet, J. W. Diclaro II. 2016. Insecticide susceptibility of natural populations of *Anopheles coluzzii* and *Anopheles gambiae* (*sensu stricto*) from Okyereko irrigation site, Ghana, West Africa. Parasit Vectors. 9: 182.

Chandre, F., F. Darriet, L. Manga, M. Akogbeto, O. Faye, J. Mouchet, P. Guillet. 1999. Status of pyrethroid resistance in *Anopheles gambiae sensu lato*. Bull. World Health Organ. 77: 230–234.

Chouaïbou, M., J. Etang, T. Brévault, P. Nwane, C. K. Hinzoumbé, R. Mimpfoundi, F. Simard. 2008. Dynamics of insecticide resistance in the malaria vector *Anopheles gambiae s.l*. from an area of extensive cotton cultivation in Northern Cameroon. Trop. Med. Inter. Health. 13: 476–486.

(CRASH) Centre de Recherche Agronomique-Savane Humide. 2012. Rapport d’activité. Centre de Recherche Agricole de la Savane Humide. 95p.

Dadzie, S. K., J. Chabi, A. Asafu-Adjaye, O. Owusu-Akrof, A. Bafoe-Wilmot, K. Malm, C. Bart-Plange, S. Coleman, M. A. Appawu, D. A. Boakye. 2017. Evaluation of piperonyl butoxide in enhancing the efficacy of pyrethroid insecticides against resistant *Anopheles gambiae s.l*. in Ghana. Malar J. 16: 342.

Diabaté, A., T. Baldet, F. Chandre, M. Akoobeto, T. R. Guiguemde, F. Darriet, C. Brengues, P. Guillet, J. Hemingway, G. J. Small, et al. 2002. The role of agricultural use of insecticides in resistance to pyrethroids in *Anopheles gambiae s.l*. in Burkina Faso. Am. J. Trop. Med. Hyg. 67: 617–622.

Djègbè, I., O. Boussari, A. Sidick, T. Martin, H. Ranson, F. Chandre, M. Akogbéto, and V. Corbel. 2011. Dynamics of insecticide resistance in malaria vectors in Benin: first evidence of the presence of L1014S *kdr* mutation in *Anopheles gambiae* from West Africa. Malar. J. 10: 261.

Djègbè, I., R. Akoton, G. Tchigossou, K. M. Ahadji-Dabla, S. M. Atoyebi, R. Adéoti, F. Zeukeng, G. K. Ketoh, and R. Djouaka. 2018. First report of the presence of L1014S Knockdown-resistance mutation in *Anopheles gambiae s.s* and *Anopheles coluzzii* from Togo, West Africa. Wellcome Open Res. 3: 30.

Elissa, N., J. Mouchet, F. Riviere, J. Y. Meunier, and K. Yao. 1993. Resistance of *Anopheles gambiae s.s.* to pyrethroids in Côte d’Ivoire. Ann. Soc. Belg. Med. Trop. 73: 291–294.

Hemingway, J., R. Shretta, T. N. C. Wells, D. Bell, A. A. Djimdé, N. Achee, G. Qi. 2016. Tools and Strategies for Malaria Control and Elimination: What Do We Need to Achieve a Grand Convergence in Malaria? PLoS Biol. 14(3): e1002380.

(IRAC) Insecticide Resistance Action Committee. 2020. http://www.irac-online.org, accessed August 17, 2020.

Ketoh, G. K., K. M. Ahadji-Dabla, J. Chabi, A. D. Amoudji, G. Y. Apetogbo, F. Awokou, I. A. Glitho. 2018. Efficacy of two PBO long lasting insecticidal nets against natural populations of *Anopheles gambiae s.l*. in experimental huts, Kolokopé, Togo. PLoS ONE. 13(7): e0192492.

Nauen R. 2007. Insecticide resistance in disease vectors of public health importance. Pest. Manag. Sci. 63: 628–633.

Pwalia, R., J. Joannides, A. Iddrisu, C. Addae, D. Acquah-Baidoo, D. Obuobi, G. Amlalo, S. Akporh, S. Gbagba, S. K. Dadzie, et al. 2019. High insecticide resistance intensity of *Anopheles gambiae* (s.l.) and low efficacy of pyrethroid LLINs in Accra, Ghana. Parasit. Vectors 12: 299.

Rowland, M., P. Boko, A. Odjo, A. Asidi, M. Akogbeto, R. N’Guessan. 2013. A new long-lasting indoor residual formulation of the organophosphate insecticide pirimiphos methyl for prolonged control of pyrethroid-resistant mosquitoes: an experimental hut trial in Benin. PLoS ONE. 8(7): e69516.

Stump, A. D., F. K. Atieli, J. M. Vulule, N. J. Besansky. 2004. Dynamics of the pyrethroid knockdown resistance allele in western Kenyan populations of *Anopheles gambiae* in response to insecticide-treated bed net trials. Am. J. Trop. Med. Hyg. 70: 591–596.

(WHO) World Health Organization. 2018. Global report on insecticide resistance in malaria vectors: 2010-2016. Global Malaria Programme. World Health Organization, Geneva, Switzerland.

(WHO) World Health Organization. 2013. Test procedures for insecticide resistance monitoring in malaria vector mosquitoes. World Health Organization, Geneva, Switzerland.

(WHO) World Health Organization. 2017. World malaria report. World Health Organization, Geneva, Switzerland.

Yadouleton, A., T. Martin, G. Padonou, F. Chandre, A. Asidi, L. Djogbenou, R. Dabiré, R. Aïkpon, M. Boko, I. Glitho, et al. 2011. Cotton pest management practices and the selection of pyrethroid resistance in *Anopheles gambiae* population in northern Benin. Parasit. Vectors 4: 60.

